# Effects of lumbar disc injury and nociception on trunk motor control during rat locomotion

**DOI:** 10.64898/2026.03.24.713869

**Authors:** Fangxin Xiao, Wendy Noort, Jia Han, Jaap H. van Dieën, Huub Maas

## Abstract

Intervertebral disc (IVD) injury is a major cause of low-back pain and can lead to structural deficits and mechanical instability. When the IVD is compromised, neuromuscular compensation by paraspinal muscles, such as the multifidus (MF) and longissimus (ML), is critical for maintaining spine stability. However, it is unknown how IVD injury and its interaction with nociception affect neuromuscular control. This study assessed the effects of IVD injury and additional muscle-derived nociception on trunk motor control during locomotion in a rat model. IVD injury was induced via needle puncture at L4/L5. One week later, hypertonic saline was injected into the lumbar MF to induce nociception. Trunk and pelvic kinematics, bilateral EMG activity of MF and ML were recorded during treadmill locomotion at baseline, one week after IVD injury, and immediately following hypertonic saline injection. Trunk and pelvic kinematics and bilateral muscle activation patterns remained largely consistent across conditions. No significant changes were found in stride duration, pelvic, lumbar and spine angle changes, variability, or movement asymmetry. MF activation was bilaterally synchronized, whereas ML showed left-right alternating activation patterns. Following IVD injury, right MF mean activation and EMG variability increased significantly compared to baseline. When muscle-derived nociception was added in the unstable spine (IVD injury) condition, left MF minimum amplitude was significantly reduced, and instability-related increases in right MF mean activation and variability were attenuated, but not fully reversed. These findings suggest that IVD injury, alone or in combination with muscle-derived nociception, elicits localized neuromuscular adaptations without disrupting the global locomotor patterns.

## Introduction

Low-back pain (LBP) is the largest cause of the world’s burden of disability, and the total number of LBP cases is predicted to increase by 36.4% worldwide by 2050 (Collaborators GBDLBP, 2023). Intervertebral disc (IVD) degeneration is one of the leading causes of LBP (Kirnaz S et al., 2022;Luoma K et al., 2000;Teraguchi M et al., 2014). Degenerated discs exhibit distinct structural and morphological alterations, such as annular fibrosus disorganization, nucleus pulposus content loss, and disc height reduction (Glaeser JD et al., 2020;Maas H et al., 2018;Rousseau MA et al., 2007). Studies have demonstrated that these structural defects can compromise mechanical properties of the spinal segment, including reduced stiffness (Gullbrand SE et al., 2017;Martin JT et al., 2013;Xiao F et al., 2025) and increased intersegmental range of motion (Gullbrand SE,Malhotra NR,Schaer TP,Zawacki Z,Martin JT,Bendigo JR,Milby AH,Dodge GR,Vresilovic EJ,Elliott DM,Mauck RL and Smith LJ, 2017;Melrose J et al., 2012).

According to the spine stability model proposed by Panjabi (Panjabi MM, 1992), the spinal stabilizing system consists of the passive subsystem (vertebrae, IVDs, ligaments), the active subsystem (paraspinal muscles and tendons), and the neural control system (peripheral and central nervous system). Dysfunction in any one of these components may trigger compensatory adaptions in the other components to maintain stability on both the short and long term (Fujii K et al., 2019;Panjabi MM, 1992). If the stability of the IVD, a component of the passive subsystem, is compromised due to injury, compensatory neuromuscular responses, such as increased back muscle activation (Stokes IA and Gardner-Morse M, 2003), may be elicited to provide active stiffness. Experimental evidence from both human and animal models have demonstrated that electrical or mechanical stimulation of sensory afferents within the IVD, ligaments, or facet joint capsule can evoke reflexive activation of the multifidus (MF) and longissimus (ML) muscle, even in the absence of structural disruption (Holm S,Indahl A and Solomonow M, 2002;Indahl A et al., 1995;Indahl A et al., 1997;Solomonow M et al., 1998).

Paraspinal muscles play a vital role in maintaining spine stability. In awake rats, lumbar MF and ML are major back extensors that are tonically active most of the time to maintain lumbar spine stability, particularly the MF (Geisler HC,Westerga J and Gramsbergen A, 1996;Wada N et al., 2006). MF connects individual lumbar vertebrae. Consequently, it can increase segmental stiffness and control intervertebral movement. ML contributes to both stabilization and movement of the trunk as a whole (Wada N,Akatani J,Miyajima N,Shimojo K and Kanda K, 2006). In addition to enhancing spine stiffness, both muscles contribute to countering lateral trunk movement caused by limb movements, with the ML being particularly involved in re-aligning the spine toward the midline (Schilling N, 2011;Wada N,Akatani J,Miyajima N,Shimojo K and Kanda K, 2006).

Structural and functional changes in MF, such as reduced cross-sectional area, fatty infiltration, and fibrosis, have been associated with LBP (Noonan AM and Brown SHM, 2021;Seyedhoseinpoor T et al., 2022). Individuals with LBP also showed altered trunk muscle activation patterns compared to healthy controls (van Dieen JH, Cholewicki J and Radebold A, 2003;van Dieen JH et al., 2019;van Dieen JH, Selen LP and Cholewicki J, 2003), and possibly related to this higher resting MF and ML stiffness (Vatovec R and Voglar M, 2024). While these changes may enhance spine stability in the short term (van Dieen JH,Cholewicki J and Radebold A, 2003) by limiting excessive motion and reducing tissue strain, they may lead to increased trunk stiffness and constrain mobility (van Dieen JH,Reeves NP,Kawchuk G,van Dillen LR and Hodges PW, 2019), and over time, may lead to elevated cumulative spinal loading, muscle remodelling (e.g., atrophy, degeneration), and further deteriorate disc health, ultimately contributing to the development or persistence of chronic LBP (Shahidi B et al., 2017;Tieppo Francio V et al., 2023;van Dieen JH,Reeves NP,Kawchuk G,van Dillen LR and Hodges PW, 2019).

Discogenic pain, consisting of nociceptive and neuropathic pain, is a distinct category of back pain (Fujii K,Yamazaki M,Kang JD,Risbud MV,Cho SK,Qureshi SA,Hecht AC and Iatridis JC, 2019). Structural IVD damage may trigger such pain, especially when accompanied by neurovascular ingrowth into the inner part of the IVD via the annular fissures (Daly C et al., 2016;Fujii K,Yamazaki M,Kang JD,Risbud MV,Cho SK,Qureshi SA,Hecht AC and Iatridis JC, 2019). However, the assessment of IVD injury related pain in animal models remains challenging. In rodents, pain is typically assessed through observation of pain-related behaviors (e.g. grooming, ‘wet-dog shakes’), measurements of functional performance (e.g. locomotion, rotarod), or responses to mechanical or thermal stimuli (Daly C,Ghosh P,Jenkin G,Oehme D and Goldschlager T, 2016). While spontaneous behavior (Olmarker K,Storkson R and Berge OG, 2002) and functional performance (Lai A et al., 2015) remained unchanged, increased hind paw sensitivity to mechanical and thermal stimuli has been consistently reported following IVD injury (Lai A,Moon A,Purmessur D,Skovrlj B,Winkelstein BA,Cho SK,Hecht AC and Iatridis JC, 2015;Mosley GE et al., 2020). Furthermore, IVD degeneration could interact with both the peripheral and central nervous systems (Fujii K,Yamazaki M,Kang JD,Risbud MV,Cho SK,Qureshi SA,Hecht AC and Iatridis JC, 2019), leading to acute inflammatory crosstalk between the IVD, dorsal root ganglion, and spinal cord with increased pain-related neuropeptide (e.g. calcitonin gene-related peptide, substance P) and neuroinflammation-related cell markers (e.g. glial fibrillary acidic protein) (Lai A et al., 2024;Orita S et al., 2011). These findings suggest that although discogenic pain may exist in rodent IVD injury models, they may not manifest in behavior and movement, yet underlying inflammatory and neurosensory changes are present and may affect neuromuscular control following IVD injury.

Previous studies have shown that nociceptive input can inhibit MF and ML muscle activation (Devecchi V et al., 2023;Xiao F et al., 2025). However, it remains unclear how IVD injury induced mechanical instability affects the neural control of these muscles, and how it interacts with nociception during locomotion. Therefore, this study aimed to assess the effects of IVD injury and its interaction with muscle-derived nociception on trunk neuromuscular control during locomotion in a rat model. We hypothesized that IVD injury will lead to increased back muscle activation and more constrained trunk motion, while the addition of muscle-derived nociception will attenuate these adaptions.

## Experimental procedures

### Study preregistration

This study was preregistered at PreclinicalTrials.eu prior to conducting the research (registration number PCTE0000367). Two interventions were preregistered, the first intervention (hypertonic saline induced muscle-derived nociception) has been reported in a previous publication (Xiao F,Noort W,Lévénez J,Han J,van Dieën JH and Maas H, 2025). Meanwhile, due to the presence of missing data, the statistical analysis method was changed from the preregistered two-way repeated measures ANOVA to paired t test.

### Animals

All experimental procedures were approval and supervised by the Animal Welfare Body at the Vrije Universiteit Amsterdam, in accordance with the Dutch law on animal research in full agreement with the Directive 2010/63/EU and approved by the Netherlands Central Commission for Animal Experiments (Permit Number AVD11200202115388).

Twelve adult male Wistar rats (*Rattus norvegicus domestica*, 330±34 gram prior to EMG implantation surgery, 9 weeks of age upon arrival) were used in this study. Only male rats were used because of the sex difference in pain response (Cairns BE et al., 2001;Capra NF and Ro JY, 2004), IVD degeneration and healing responses after IVD injury (Mosley GE et al., 2019).

### Experimental protocol

This study was conducted using the same cohort of rats as in a prior experiment involving intramuscular hypertonic saline (HS) injection (Xiao F,Noort W,Lévénez J,Han J,van Dieën JH and Maas H, 2025). The surgical procedures for EMG electrode implantation, treadmill locomotion training, intramuscular hypertonic saline injection, in vivo measurement, and IVD injury protocols were identical to those previously reported (Xiao F,Noort W,Lévénez J,Han J,van Dieën JH and Maas H, 2025) (Xiao F,Noort W,Lévénez J,Han J,van Dieën JH and Maas H, 2025), and are briefly summarized here.

The timeline of the experimental design is shown in Fig.1. Following one week of acclimatization, the rats underwent two weeks of treadmill locomotion training (Exer 3/6, Columbus Instruments). Subsequently, EMG electrodes were implanted bilaterally into the multifidus (MF) and medial longissimus (ML) muscles at the L4-L5 vertebral level. Kinematic markers were placed on L2 and S1 spinous processes and bilateral iliac crests. After two-weeks recovery, in vivo measurements were performed to collect locomotion videos and EMG signals while the rats trotted on the treadmill at a fixed speed of 0.5 m/s. Locomotion videos (200 frames/second) and EMG signals (sampling frequency: 3123 Hz) were synchronized by an electronic trigger pulse to the controller.

**Fig. 1.**
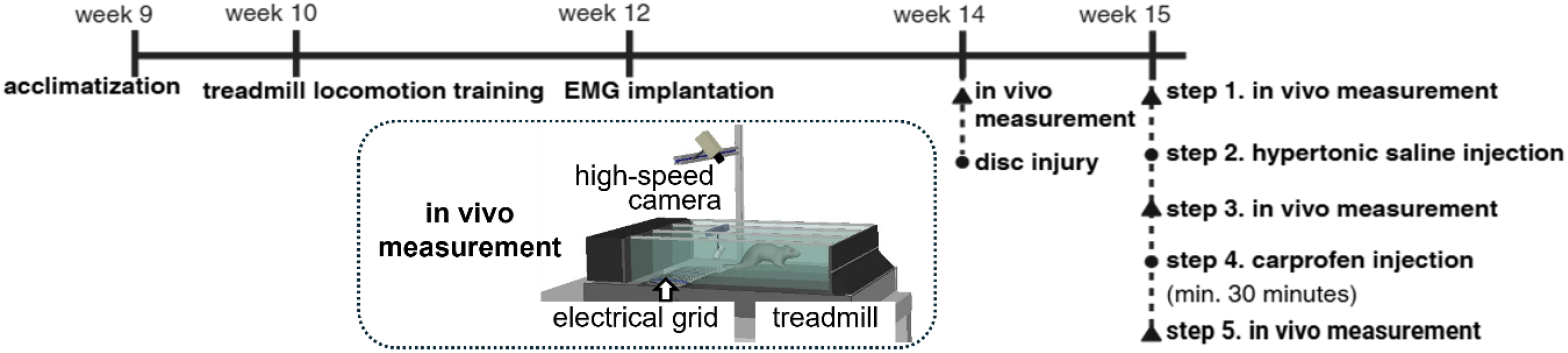
Overview of experimental procedures and timeline (week: age of the rats).

After the baseline recording, the L4/5 IVD was injured to induce local spinal instability. The IVD injury was created by puncturing the IVD with a 21-gauge needle to a depth of 2.5mm, fully penetrating the nucleus pulposus (Xiao F,Noort W,Lévénez J,Han J,van Dieën JH and Maas H, 2025). One week post IVD injury, in vivo measurements were repeated to evaluate locomotion and muscular responses associated with the IVD injury. To assess the interaction between IVD injury and muscle-derived nociception (condition: HS), a subsequent intervention was performed in which 100µl hypertonic saline (5.8%) was injected into either the left or right MF at L4-L5 vertebral level. Immediately after this injection, another round of in vivo measurements was conducted. Finally, to assess if motor control was affected by pain at one week after IVD injury, an analgesic condition was included as an additional control, in which carprofen (3 mg/kg, Rimadyl®, Zoetis B.V., Capelle a/d Ijssel, The Netherlands) was administrated subcutaneously to the rats at least 30 minutes prior to the final in vivo measurement.

The rats were housed in pairs, except that after the EMG electrode implantation surgery, the two rats in the same cage were separated by a cage divider. All rats had *ad libitum* access to food and water and were housed under a 12-h light/dark cycle.

### Data analysis

Kinematic and EMG data analysis followed the same procedures as described previously (Xiao F,Noort W,Lévénez J,Han J,van Dieën JH and Maas H, 2025). Briefly, the locomotion videos were analyzed in Deeplabcut (Nath T et al., 2019) to obtain the time series of the lumbar and pelvic angles, which were defined as the angles between the markers L2-S1, left-right iliac crest, and the horizontal line (positive direction: right), respectively. Segmental angle outliers (pelvic angles outside 250-300 degree and lumbar angles outside 0-30 degree and 330-360 degree) were excluded and interpolated (Piecewise Cubic Hermite Interpolating Polynomial). The resulting data were low-pass filtered (5 Hz, 3^rd^ order zero-lag Butterworth).

Stride cycles were identified using the pelvic angle, with each cycle starting at its minimum value. Only trotting strides at constant speed (cycle duration: 0.2-0.4 s) were included, galloping strides and those with excessive forward-backward acceleration or lateral motion were excluded. Angle and EMG data were time-normalized to 100 data points per stride cycle. For each rat, data were averaged across stride cycles from the same measurement session, EMG envelope amplitude was further normalized to the maximum value of the mean EMG recorded during baseline.

Changes in lumbar and pelvic angles were calculated based on time-normalized data. To correct for potential asymmetry caused by marker placement, the mean of the time series was subtracted. The spine angle was calculated as the difference between lumbar and pelvic angles. Timing of peak pelvic angle was assessed. Kinematic variability was quantified as the mean standard deviation across stride cycles at each normalized time point. Movement asymmetry was calculated as the standard deviation of the differences between mirrored normalized time points in the first and second halves of the spine angle curve, divided by half of the peak-to-peak amplitude. EMG parameters were peak, minimum, and average amplitudes of the amplitude-normalized EMG envelope. EMG variability was calculated in the same manner as for kinematics. Pearson correlation coefficients were calculated between the left and right sides of each muscle based on the amplitude-normalized EMG envelopes within each condition. All parameters were averaged within rats.

### Statistical analysis

EMG data from specific channels were excluded when there was a bad signal-to-noise ratio, electrode malfunction or inappropriate placement of the electrodes. The spm1d-package for n-dimensional statistical parametric mapping (SPM) (Pataky TC, 2010) was used to compare the time series of kinematic and EMG activity data between different conditions. To explore whether potential pain from IVD injury elicited measurable changes in neuromuscular control, *spm1d*.*stats*.*ttest_paired* was conducted to compare the time series of kinematic and EMG activity data between the IVD injury and analgesia conditions. Additionally, paired t-tests were used to compare the outcome parameters from these two conditions including both kinematics (pelvic/lumbar/spine angle change and variability, movement asymmetry) and EMG activity (peak amplitude, minimum amplitude, mean amplitude, variability, left-right correlation). Since no significant differences were detected, data from the IVD injury condition were used for subsequent statistical analyses.

To assess the effects of spine instability and its interaction with muscle-derived nociception on kinematics and EMG activity time series data, *spm1d*.*stats*.*ttest_paired* was used to assess differences in the mean and variability between (1) baseline and IVD injury (or analgesia, depending on above paired t-test results), and (2) IVD injury and IVD injury combined with nociception (condition: HS). To assess the effects of condition on each kinematic and EMG outcome parameter, separate paired t-tests were performed to evaluate differences between (1) baseline and IVD injury and (2) IVD injury and HS conditions. Left-right coordination within each muscle was assessed by calculating the Pearson correlation coefficients between the amplitude normalized EMG time series of the left and right sides (MF and ML separately) for each condition. Correlation coefficients were Fisher z-transformed prior to statistical comparison across conditions using paired t-tests. Significance level was set at 0.05. MATLAB R2021a (MathWorks, Inc., Natick, MA, United States) was used for SPM analysis. Jamovi (2.3.28) was used for paired t-tests. Normality of paired differences was assessed using Shapiro-Wilk tests, no substantial deviations from normality were detected.

## Results

None of rats showed pain-related behaviors (e.g., increased grooming, ‘wet-dog shakes’) post IVD injury until the end of experiments. During the IVD injury recovery period, several rats had chewed the skin incisions and sutures open. Local wounds were treated and re-sutured immediately after observation. Data from one rat were excluded because of poor pelvic marker recognition and malfunction of EMG electrodes. Some EMG data were excluded from specific muscles (left MF, n=4; right MF, n=2; left ML, n=1), because of a low signal-to-noise ratio, improper placement or malfunctioning of electrodes. In addition, one rat in each of the IVD injury and HS conditions was excluded due to the lack of stable trotting gait during data collection. In the HS condition, EMG signals were recorded at 2.6±1.4 min (range: 1-5 min) after hypertonic saline injection.

As no significant differences in kinematics and EMG activity were found between the IVD injury and analgesia conditions in either SPM analysis (Fig.S1-S2) or paired t-tests (Table 1-3), data from the IVD injury condition were used to represent the isolated effects of IVD injury.

**Table 1.**
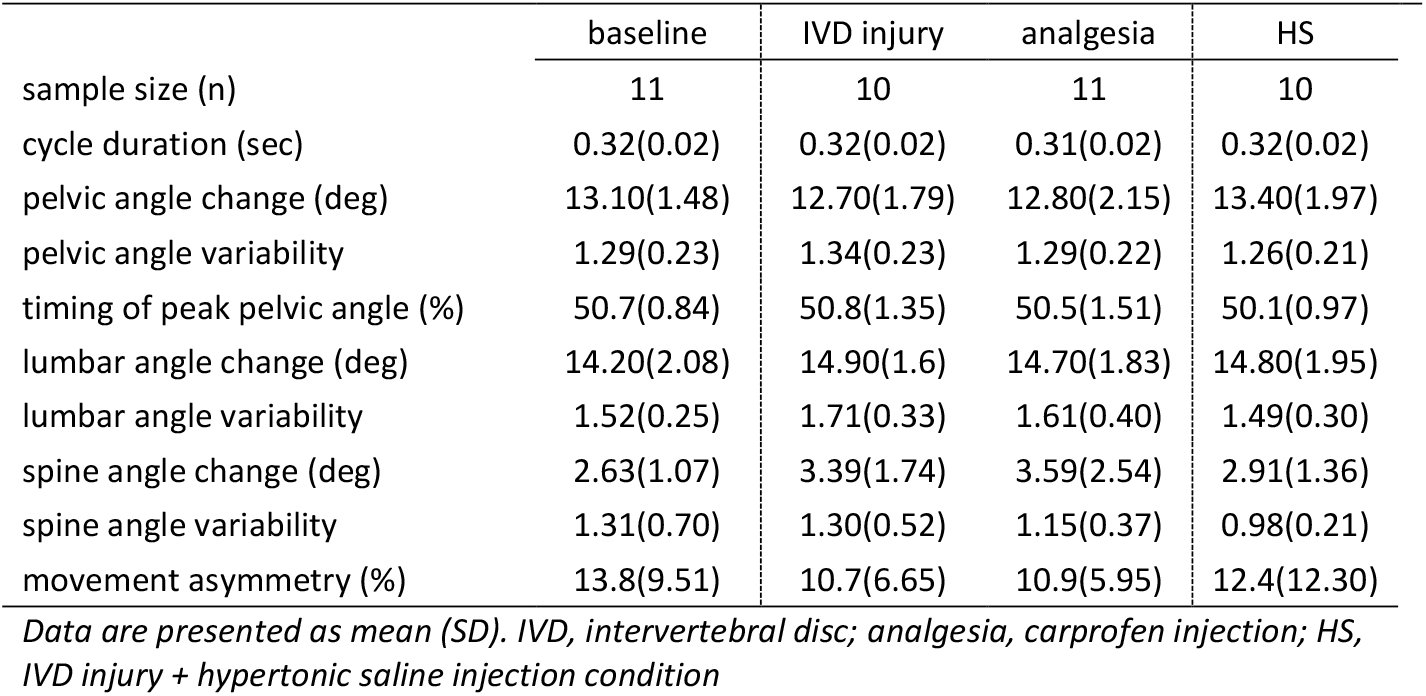
Kinematic outcomes across conditions.

**Table 2.** EMG outcomes across conditions.

|  |  | baseline |  | IVD injury |  | analgesia |  | HS |  |
| --- | --- | --- | --- | --- | --- | --- | --- | --- | --- |
|  |  | right | left | right | left | right | left | right | left |
| **MF** | n | 9 | 7 | 8 | 6 | 9 | 7 | 8 | 6 |
|  | peak amplitude | 1.00 | 1.00 | 1.37<br>(0.51) | 1.21<br>(0.72) | 1.19<br>(0.48) | 1.32<br>(1.11) | 1.18<br>(0.72) | 1.32<br>(0.82) |
|  | min amplitude | 0.23<br>(0.14) | 0.32<br>(0.18) | 0.30<br>(0.26) | 0.21<br>(0.24) | 0.24<br>(0.23) | 0.29<br>(0.19) | 0.28<br>(0.26) | 0.36<br>(0.33) |
|  | mean amplitude | 0.49<br>(0.12) | 0.57<br>(0.15) | 0.65<br>(0.27) | 0.63<br>(0.41) | 0.55<br>(0.31) | 0.62<br>(0.43) | 0.54<br>(0.35) | 0.71<br>(0.50) |
|  | variability | 0.27<br>(0.06) | 0.33<br>(0.08) | 0.41<br>(0.13) | 0.43<br>(0.28) | 0.33<br>(0.17) | 0.45<br>(0.38) | 0.30<br>(0.12) | 0.46<br>(0.40) |
| **ML** | n | 6 | 6 | 5 | 5 | 6 | 6 | 5 | 5 |
|  | peak amplitude | 1.00 | 1.00 | 0.89<br>(0.48) | 0.86<br>(0.36) | 0.85<br>(0.44) | 0.78<br>(0.21) | 0.82<br>(0.27) | 0.95<br>(0.27) |
|  | min amplitude | 0.39<br>(0.12) | 0.38<br>(0.22) | 0.29<br>(0.06) | 0.30<br>(0.13) | 0.30<br>(0.05) | 0.26<br>(0.09) | 0.34<br>(0.05) | 0.33<br>(0.12) |
|  | mean amplitude | 0.68<br>(0.10) | 0.70<br>(0.12) | 0.58<br>(0.25) | 0.58<br>(0.24) | 0.55<br>(0.24) | 0.50<br>(0.17) | 0.56<br>(0.16) | 0.63<br>(0.19) |
|  | variability | 0.32<br>(0.08) | 0.37<br>(0.15) | 0.27<br>(0.15) | 0.32<br>(0.19) | 0.24<br>(0.15) | 0.25<br>(0.14) | 0.28<br>(0.15) | 0.35<br>(0.12) |
| **Pearson correlation** |  |  |  |  |  |  |  |  |  |
| **MF** | n | 6 |  | 5 |  | 6 |  | 5 |  |
|  | L-R correlation | 0.78(0.13) |  | 0.51(0.61) |  | 0.59(0.44) |  | 0.55(0.52) |  |
| **ML** | n | 6 |  | 5 |  | 6 |  | 5 |  |
|  | L-R correlation | -0.37(0.26) |  | -0.33(0.44) |  | -0.16(0.33) |  | -0.42(0.41) |  |

Data are presented as mean(SD). n, sample size; IVD, intervertebral disc; analgesia, carprofen injection; HS, IVD injury + hypertonic saline injection condition; MF, multifidus muscle; ML: medial longissimus muscle; L-R correlation, Pearson correlation coefficient between left and right EMG time series of the same muscle.

**Table 3.** Paired t-test results for kinematic and EMG outcomes between IVD injury and analgesia conditions.

|  |  | t-value | df | p | 95% CI |  | effect size |
| --- | --- | --- | --- | --- | --- | --- | --- |
|  |  |  |  |  | lower | upper |  |
| **kinematic** | cycle duration | 0.62 | 9 | 0.548 | -0.009 | 0.015 | 0.20 |
|  | pelvic angle change | -0.07 | 9 | 0.948 | -1.011 | 0.953 | -0.02 |
|  | pelvic angle variability | 0.51 | 9 | 0.619 | -0.082 | 0.130 | 0.16 |
|  | timing peak pelvic angle | 1.04 | 9 | 0.327 | -0.333 | 0.897 | 0.33 |
|  | lumbar angle change | 0.34 | 9 | 0.740 | -1.004 | 1.362 | 0.11 |
|  | lumbar angle variability | 0.41 | 9 | 0.693 | -0.227 | 0.327 | 0.13 |
|  | spine angle change | -0.92 | 9 | 0.383 | -1.374 | 0.582 | -0.29 |
|  | spine angle variability | 0.63 | 9 | 0.545 | -0.271 | 0.480 | 0.20 |
|  | movement asymmetry | -0.50 | 9 | 0.632 | -5.925 | 3.795 | -0.16 |
| **EMG** |  |  |  |  |  |  |  |
| peak amplitude | MF right | 0.82 | 7 | 0.441 | -0.154 | 0.317 | 0.29 |
|  | MF left | 1.49 | 5 | 0.196 | -0.178 | 0.671 | 0.61 |
|  | ML right | 0.06 | 4 | 0.955 | -0.117 | 0.122 | 0.03 |
|  | ML left | 0.83 | 4 | 0.452 | -0.230 | 0.427 | 0.37 |
| minimum amplitude | MF right | 0.67 | 7 | 0.524 | -0.091 | 0.164 | 0.24 |
|  | MF left | 0.43 | 5 | 0.688 | -0.106 | 0.148 | 0.17 |
|  | ML right | -0.35 | 4 | 0.743 | -0.085 | 0.066 | -0.16 |
|  | ML left | 1.68 | 4 | 0.168 | -0.024 | 0.097 | 0.75 |
| mean amplitude | MF right | 1.24 | 7 | 0.254 | -0.055 | 0.176 | 0.44 |
|  | MF left | 1.58 | 5 | 0.174 | -0.074 | 0.313 | 0.65 |
|  | ML right | 0.79 | 4 | 0.472 | -0.066 | 0.118 | 0.35 |
|  | ML left | 1.72 | 4 | 0.160 | -0.047 | 0.201 | 0.77 |
| variability | MF right | 1.19 | 7 | 0.272 | -0.058 | 0.177 | 0.42 |
|  | MF left | 1.22 | 5 | 0.276 | -0.096 | 0.270 | 0.50 |
|  | ML right | 1.41 | 4 | 0.232 | -0.033 | 0.099 | 0.63 |
|  | ML left | 1.39 | 4 | 0.237 | -0.081 | 0.244 | 0.62 |
| L-R correlation | MF | 0.01 | 4 | 0.989 | -0.871 | 0.882 | 0.01 |
|  | ML | -0.21 | 4 | 0.842 | -0.968 | 0.789 | -0.10 |

IVD, intervertebral disc; MF, multifidus muscle; ML, medial longissimus muscle; df, degree of freedom; CI, confidence interval; effect size, Cohen's D; L-R correlation, Pearson correlation coefficient between left and right amplitude normalized EMG time series of the same muscle, correlation coefficients were Fisher z-transformed for statistical analysis. \*significant difference

Time series of the pelvic, lumbar, and spine angles exhibited highly consistent patterns over the stride cycle across the baseline, IVD injury, HS conditions (Fig.2), no significant differences in the curves of the angular means (Fig.S3-4) and variabilities (Fig.S5-6) were identified in the SPM analysis between (1) baseline and IVD injury and (2) IVD injury and HS conditions (all *p*>0.05). The pelvic and lumbar angles peaked approximately at the middle (50%) of the stride cycle. The spine angle followed a sinusoidal curve pattern (Fig.2C), with minor, non-significant deviations in the timing and amplitude across conditions. Paired t-tests revealed no significant differences between baseline and IVD injury for any of the kinematic parameters (Table 4). Similarly, no significant differences were observed between IVD injury and HS for cycle duration, segmental (pelvic, lumbar, and spine) angle changes and variability, or movement asymmetry (Table 5). Although SPM analysis did not reveal significant differences in the continuous angular waveform, paired t-test showed a modest but statistically significant earlier timing of peak pelvic angle in the HS condition compared to IVD injury (mean difference: 0.7% of the stride cycle, *p*=0.038, t=2.48, effect size=0.83). Both left and right MF showed a distinct double-peaked activation pattern aligned with the stride cycle (Fig.3A-B). These peaks were temporally aligned with changes in pelvic and lumbar angles, particularly with the timing of the minimum and maximum angles, corresponding to the stance phase of the left (start/end-stride) and right (mid-stride) hind paw. Bilateral MF activation patterns were preserved across conditions (baseline, IVD injury, HS) with similar timing and shape. SPM analysis revealed no significant differences in the EMG time series for either the mean (Fig.S7-8, A-B) or the stride-to-stride variability (Fig.S9-10, A-B) between (1) baseline and IVD injury and (2) IVD injury and HS conditions (all *p*>0.05). Pearson correlation coefficients between left and right MF EMG curves were 0.78 at baseline, 0.51 following IVD injury, and 0.55 in HS condition, indicating moderate to strong bilateral MF synchronized activation across conditions. Paired t-tests revealed no significant differences between baseline and IVD injury, nor between IVD injury and HS, in the correlation coefficients (*p*>0.05), suggesting that neither IVD injury nor its combination with muscle-derived nociception significantly altered this bilateral MF coordination.

**Fig. 2.**
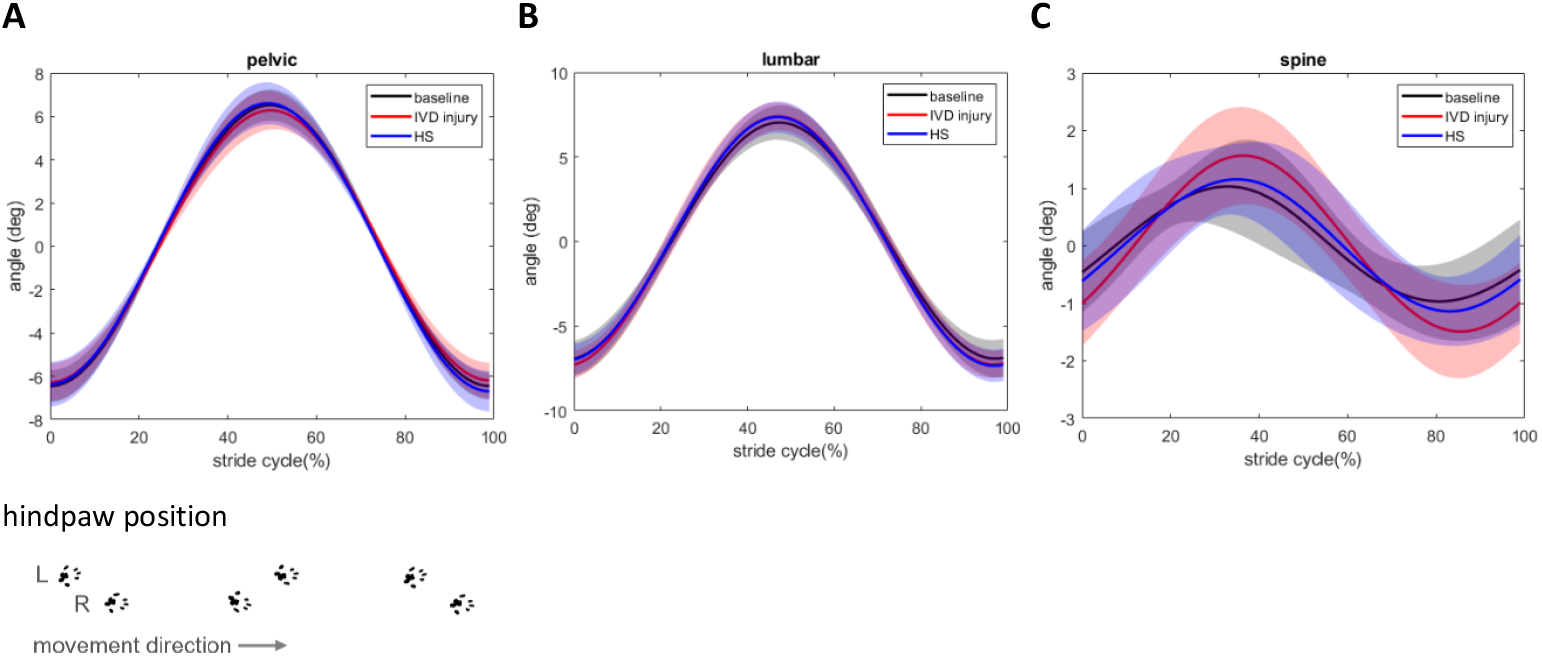
Joint angle changes during locomotion. **(A)** pelvic, **(B)** lumbar, **(c)** spine. Joint angle data were averaged cross all rats and normalized to stride cycle duration. Shaded area represents mean±1SD. Treadmill speed was at 0.5m/s at all conditions. IVD, intervertebral disc; HS, IVD injury + hypertonic saline injection condition.

**Fig. 3.**
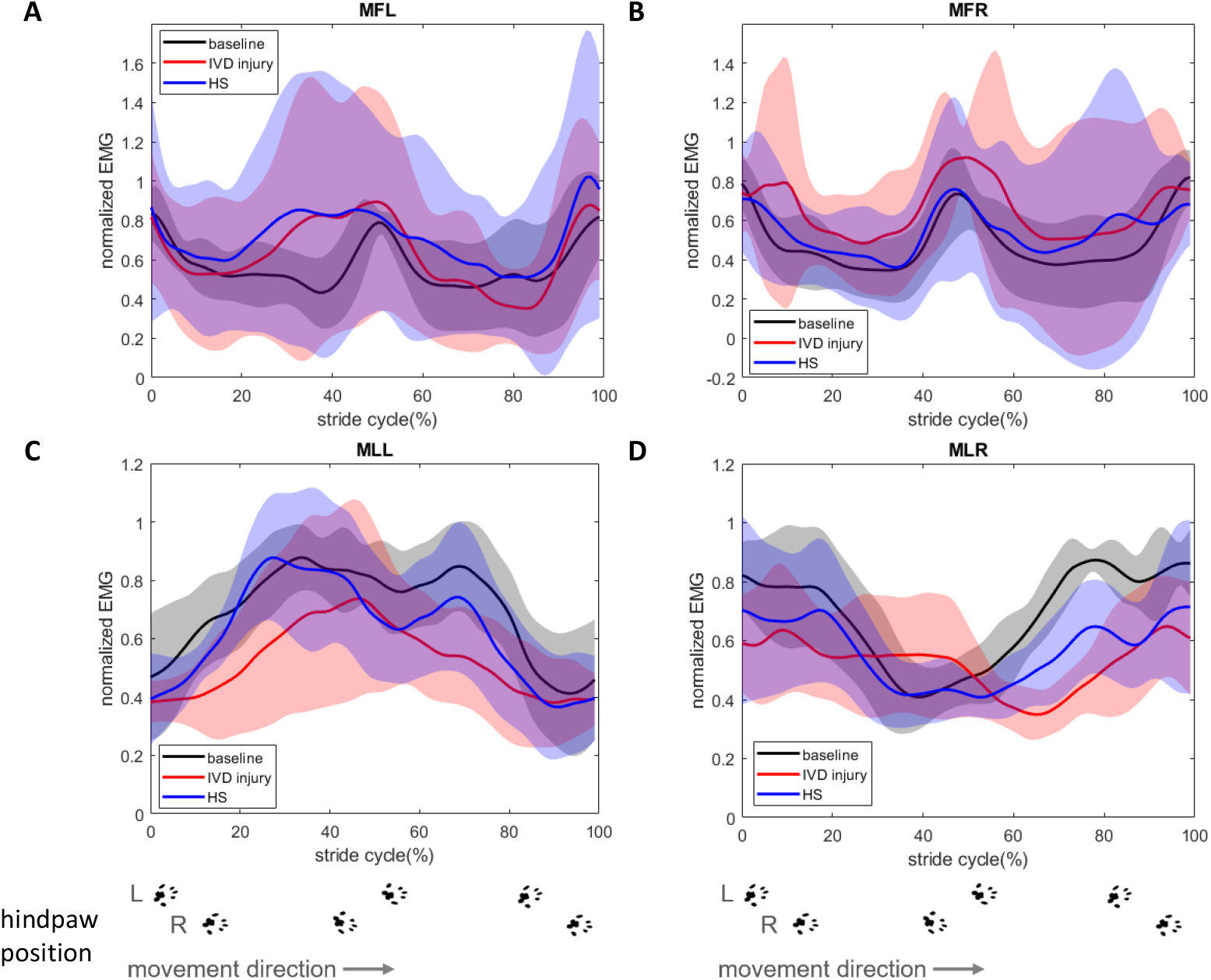
Filtered rectified EMG envelope of **(A)** left (MFL) and **(B)** right (MFR) multifidus muscle, **(C)** left (MLL) and **(D)** right (MLR) longissimus muscle during locomotion. EMG data were normalized to the stride cycle duration and peak amplitude measured during baseline. Shaded area represents mean±1SD. Treadmill speed was at 0.5m/s at all conditions. IVD, intervertebral disc; HS, IVD injury + hypertonic saline injection condition.

**Table 4.** Paired t-test results for kinematic and EMG outcomes between baseline and IVD injury conditions.

|  |  | t-value | df | p | 95% CI |  | effect size |
| --- | --- | --- | --- | --- | --- | --- | --- |
|  |  |  |  |  | lower | upper |  |
| **kinematic** | cycle duration | 1.17 | 9 | 0.271 | -0.281 | 1.004 | 0.37 |
|  | pelvic angle change | 1.37 | 9 | 0.204 | -0.228 | 1.073 | 0.43 |
|  | pelvic angle variability | -0.45 | 9 | 0.666 | -0.760 | 0.486 | -0.14 |
|  | timing peak pelvic angle | -0.11 | 9 | 0.916 | -0.653 | 0.587 | -0.03 |
|  | lumbar angle change | -0.99 | 9 | 0.347 | -0.942 | 0.330 | -0.31 |
|  | lumbar angle variability | -1.68 | 9 | 0.128 | -1.182 | 0.148 | -0.53 |
|  | spine angle change | -1.00 | 9 | 0.345 | -0.943 | 0.329 | -0.32 |
|  | spine angle variability | 0.66 | 9 | 0.525 | -0.424 | 0.831 | 0.21 |
|  | movement asymmetry | 0.90 | 9 | 0.392 | -0.356 | 0.910 | 0.28 |
| **EMG** |  |  |  |  |  |  |  |
| peak amplitude | MF right | -1.80 | 7 | 0.114 | -1.386 | 0.147 | -0.64 |
|  | MF left | -0.74 | 4 | 0.502 | -1.215 | 0.593 | -0.33 |
|  | ML right | 0.52 | 4 | 0.633 | -0.673 | 1.107 | 0.23 |
|  | ML left | 0.85 | 4 | 0.442 | -0.552 | 1.273 | 0.38 |
| minimum amplitude | MF right | -0.89 | 7 | 0.403 | -1.016 | 0.407 | -0.32 |
|  | MF left | -0.84 | 4 | 0.450 | -1.265 | 0.558 | -0.37 |
|  | ML right | 1.33 | 4 | 0.255 | -0.397 | 1.528 | 0.59 |
|  | ML left | 1.58 | 4 | 0.188 | -0.320 | 1.673 | 0.71 |
| mean amplitude | MF right | -2.50 | 7 | **0.041\*** | -1.691 | -0.034 | -0.88 |
|  | MF left | -0.71 | 4 | 0.517 | -1.201 | 0.603 | -0.32 |
|  | ML right | 0.83 | 4 | 0.455 | -0.562 | 1.260 | 0.37 |
|  | ML left | 1.70 | 4 | 0.164 | -0.285 | 1.742 | 0.76 |
| variability | MF right | -3.35 | 6 | **0.015\*** | -2.261 | -0.223 | -1.27 |
|  | MF left | -0.90 | 4 | 0.418 | -1.299 | 0.535 | -0.40 |
|  | ML right | 0.63 | 4 | 0.560 | -0.629 | 1.164 | 0.28 |
|  | ML left | 0.66 | 4 | 0.544 | -0.619 | 1.178 | 0.30 |
| L-R correlation | MF | 1.08 | 4 | 0.339 | -0.475 | 1.394 | 0.48 |
|  | ML | 1.12 | 3 | 0.344 | -0.545 | 1.593 | 0.56 |

IVD, intervertebral disc; MF, multifidus muscle; ML, medial longissimus muscle; df, degree of freedom; CI, confidence interval; effect size, Cohen's D; L-R correlation, Pearson correlation coefficient between left and right amplitude normalized EMG time series of the same muscle, correlation coefficients were Fisher z-transformed for statistical analysis. \*significant difference

**Table 5.**
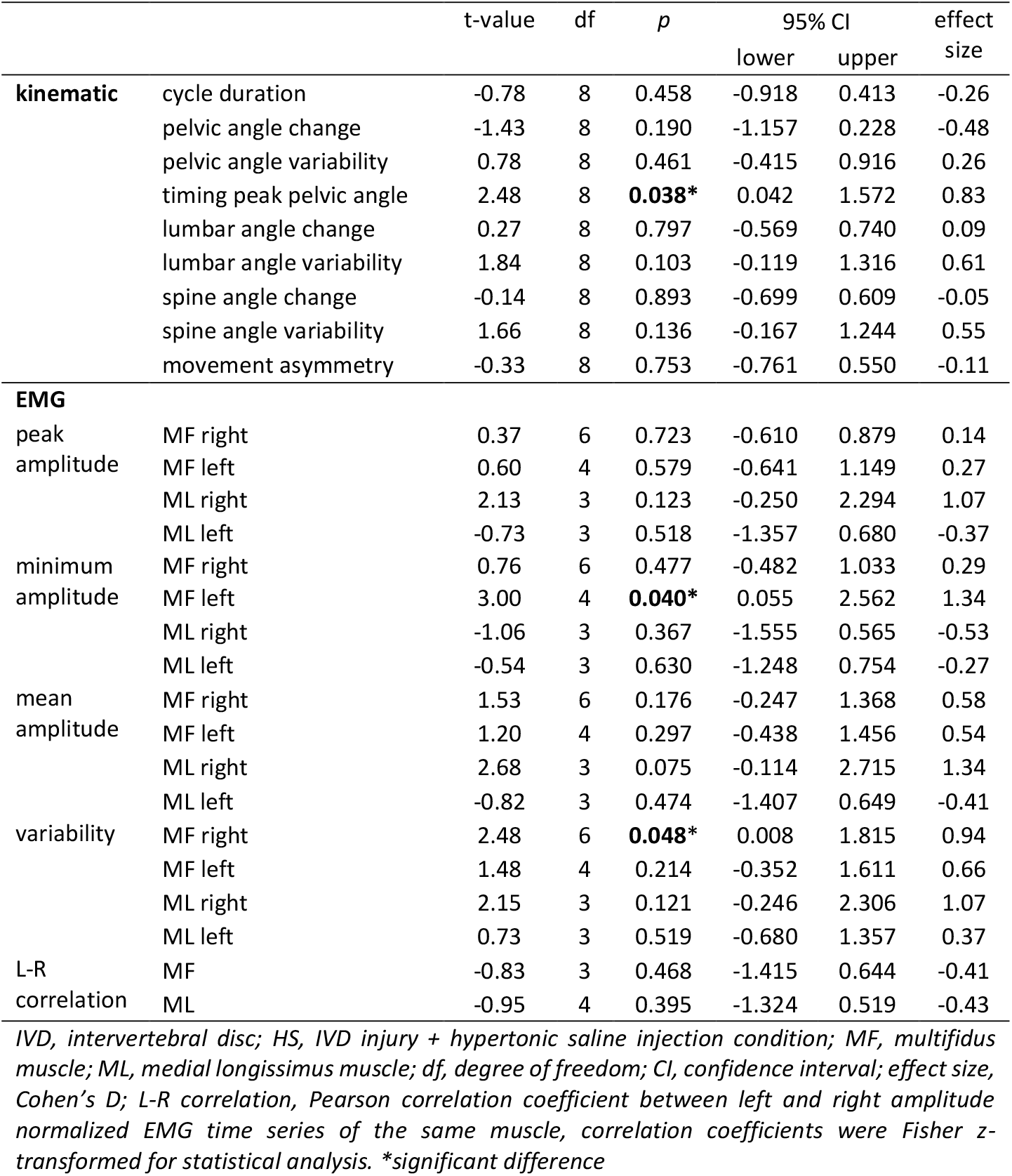
Paired t-test results for kinematic and EMG outcomes between IVD injury and HS conditions.

Overall EMG amplitudes appeared elevated in the IVD injury condition. Paired t-tests revealed that the mean amplitude of only the right MF was significantly higher in the IVD injury condition compared to baseline (*p*=0.041, t=-2.50 effect size=-0.88), while no significant difference was observed between IVD injury and HS (*p*=0.176). No significant differences were observed for peak or minimum amplitudes of the right MF (Fig.4, Table 5). Stride-to-stride variability in right MF activation followed the time course of the mean EMG curve, with greater variability during the main burst phases of the stride cycle (Fig.S6B). For the mean variability across the stride, right MF EMG variability was significantly higher following IVD injury compared to baseline (*p*=0.015, t=-3.35, effect size=-1.27), and the variability was significantly reduced in the HS condition compared to IVD injury (*p*=0.048, t=2.48, effect size=0.94). For the left MF, minimum amplitude did not differ between baseline and IVD injury (*p*=0.403), but was significantly lower in the HS condition compared to IVD injury (*p*=0.040, t=3.00, effect size=1.34). No other significant differences were observed for left MF activation parameters.

**Fig. 4.**
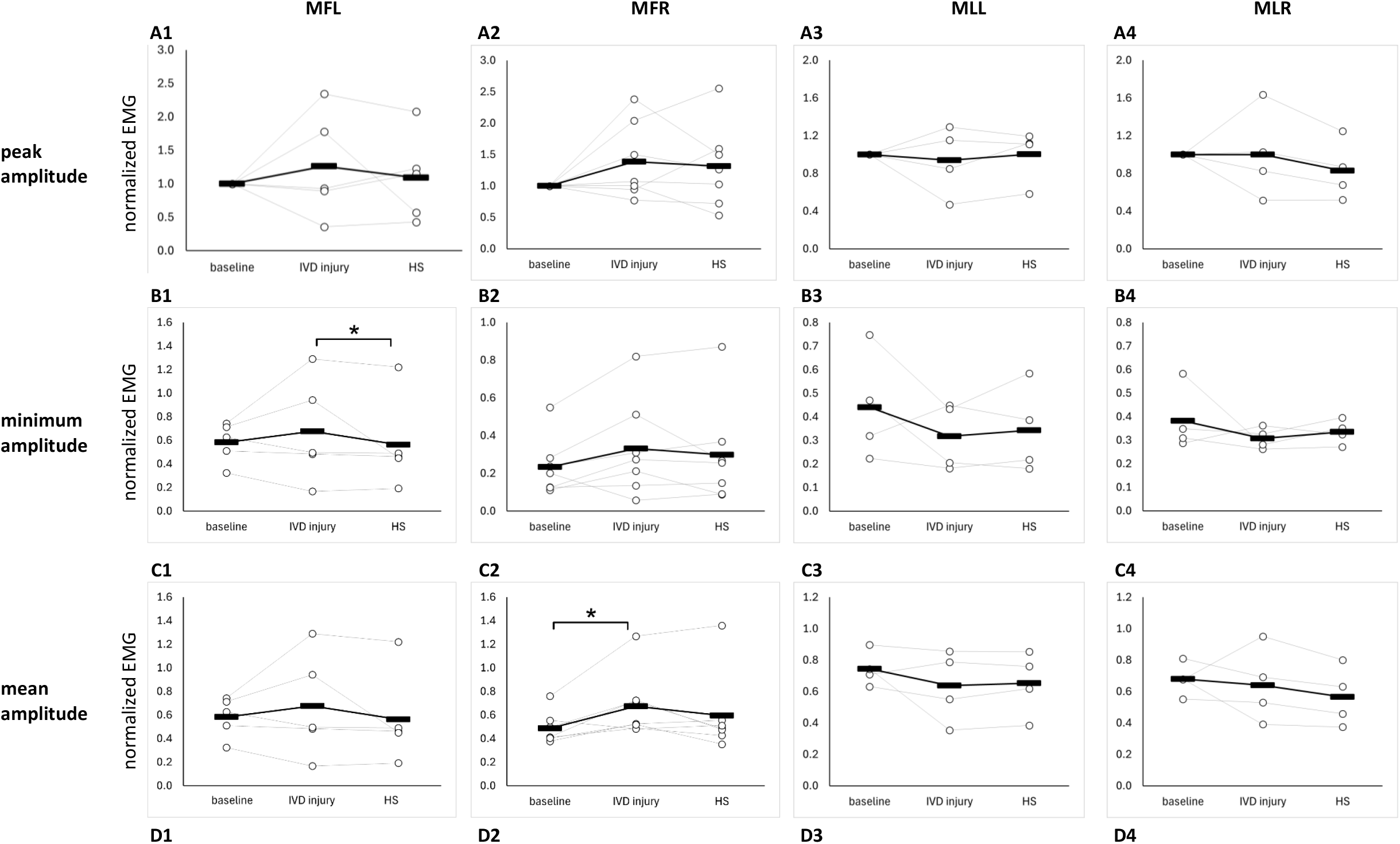
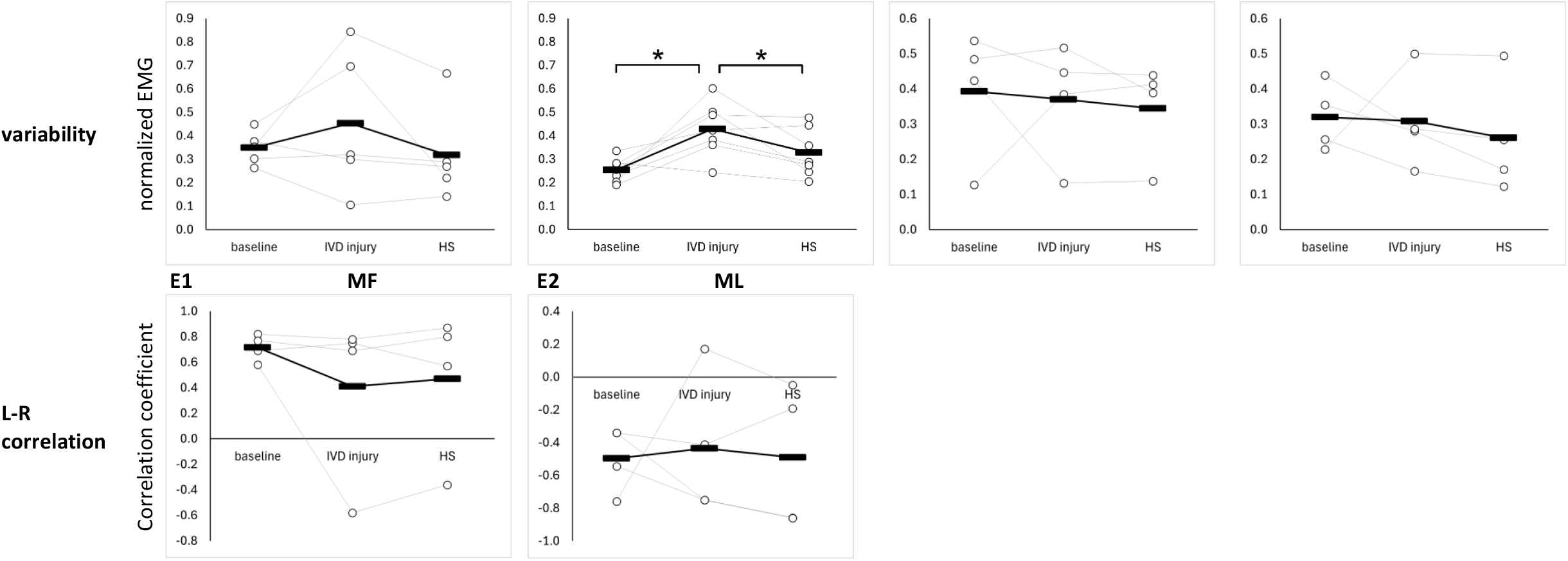
Normalized EMG outcomes during locomotion for all conditions. **(A)** peak amplitude, **(B)** minimum amplitude, **(C)** mean amplitude, **(D)** variability, **(E)** L-R correlation. Hollow circles represent individual data of each rat, black bars represents the group mean. MF, multifidus muscle; ML, medial longissimus muscle; L, left; R, right; IVD, intervertebral disc; HS, IVD injury + hypertonic saline injection condition; \**p*<0.05, significant difference.

As in our previous paper (Xiao F,Noort W,Lévénez J,Han J,van Dieën JH and Maas H, 2025), data analysis for the ML was restricted to those rats (n=6) whose baseline ML EMG patterns matched with established descriptions in literature(Geisler HC,Westerga J and Gramsbergen A, 1996). Both left and right ML exhibited a single, broad activation pattern during the stride cycle across all conditions (baseline, IVD injury, HS) (Fig.3C-D). There was alternating activation between left and right ML, with increased activation corresponding to the swing phase of the ipsilateral hind paw. Moderate negative Pearson correlation coefficients between left and right ML (baseline: −0.37, IVD injury: −0.33, HS: −0.42) indicate alternating activation patterns between the two sides in all conditions. A slight downward shift in the EMG envelopes was observed in the IVD injury condition compared to baseline, however, SPM analysis revealed no significant differences between baseline and IVD injury (Fig.S7C-D). Likewise, no significant differences were observed between IVD injury and HS conditions in the ML EMG time series data (Fig.S8C-D). Similarly, SPM analysis revealed no significant differences in the ML EMG variability time series data (Fig.S9-10, C-D). Consistent with the SPM results, paired t-tests for ML EMG outcomes showed no significant differences between baseline and IVD injury, nor between IVD injury and HS, for peak, minimum, or mean amplitude, variability, or left-right correlation coefficients (Fig.4, Table 5).

## Discussion

This study investigated the effects of IVD injury and its interaction with muscle-derived nociception on trunk neuromuscular control during locomotion in a rat model. The main findings indicate that neither IVD injury alone nor in combination with muscle-derived nociception substantially altered spine and pelvis kinematics. Trunk muscle activation patterns were largely preserved across conditions. Following IVD injury, localized increases of EMG mean amplitude and variability were observed in the right MF, when muscle-derived nociception was added in the IVD injury condition, EMG minimum amplitude and variability were significantly reduced both in the left and right MF. The increases of EMG mean amplitude and variability in the MF after IVD injury were attenuated with nociception, but not fully reversed.

## Nociception arising from IVD injury did not alter motor output

No significant differences were found between the IVD injury and analgesia (carprofen injection) conditions in either kinematic or EMG outcomes. Consistent with Lai et al. (Lai A,Moon A,Purmessur D,Skovrlj B,Winkelstein BA,Cho SK,Hecht AC and Iatridis JC, 2015), there were no changes in pain-related behaviors or functional performance (e.g. rotarod, inclined plane) following IVD injury, despite increased hind paw mechanical and thermal sensitivities. This suggests that nociceptive input arising from the injured disc was either minimal or insufficient to alter motor output in this model.

## Kinematic consistency during IVD injury and additional muscle-derived nociception

Although we hypothesized that IVD injury would constrain segmental motion, the observed kinematic differences were small and not significant. Additional nociceptive input through hypertonic saline injection into lumbar MF also did not significantly change the trunk and pelvic angle changes, variability, cycle duration, or movement asymmetry relative to isolated IVD injury. However, a modest but statistically significant earlier timing of peak pelvic angle (0.7% of the stride cycle) was observed in the HS condition compared to IVD injury. In the absence of amplitude or variability changes, this temporal shift likely reflects subtle coordination adjustments rather than alteration of the global motor output.

These findings align with open field test results, showing no significant gait abnormalities in rats with IVD injury, including distance traveled, stance time imbalance, temporal symmetry, spatial symmetry, step width, stride length, or duty factor (Lee FS et al., 2024). Our previous muscle-derived nociception-only study also did not find significant kinematic effects (Xiao F,Noort W,Lévénez J,Han J,van Dieën JH and Maas H, 2025). Similarly, human studies also reported no significant differences in the amplitude of motion in lumbar spine, pelvis, or hip between individuals with and without LBP (Smith JA et al., 2022). However, detailed gait analysis in rats using a computer-assisted gait analysis system with paw print and timing detection found longer stance phases and shorter strides at one to two weeks following IVD injury (Miyagi M et al., 2013). In a facetectomy induced spine instability rat model, no gait abnormalities were observed in the first six weeks post injury, but a reduced swing speed was observed at week seven (Fukui D et al., 2015). With our experimental setup, it was not possible to analyze stance and swing phases separately, as locomotion was recorded from the back of the rats. Further studies involving more in-depth gait analysis and a longer observational period may be necessary to detect potential gait adaptations or impairments.

The absence of alterations in spine angle may be explained by the limited effects of IVD injury on segmental mechanical properties. Mechanical testing of the spinal segments from the same model revealed that IVD injury did reduce peak stiffness and peak passive moments in flexion, but lateral bending mechanics remained unaffected (Xiao F,Noort W,Lévénez J,Han J,van Dieën JH and Maas H, 2025). These results suggest that neither IVD injury nor the additional muscle-derived nociception were sufficient to disrupt overall trunk and pelvis kinematics during locomotion. Future studies could assess effects of IVD injury on sagittal plane spine movement, such as using high-speed X-ray video, which could eliminate artifacts caused by soft tissue movement (Bauman JM and Chang YH, 2010).

## Localized neuromuscular adaptations to IVD injury and additional muscle-derived nociception

Muscle activation patterns were preserved across muscles and conditions, suggesting that overall trunk neuromuscular control remained stable following IVD injury and additional muscle-derived nociceptive input. Localized neuromuscular changes were observed in the MF. IVD injury resulted in an increase in EMG mean amplitude and variability of the right MF compared to baseline. These findings are consistent with the hypothesis that reduced passive spinal stability elicits increased activation of segmentally inserting lumbar muscles to enhance mechanical stability. According to the spine stability framework proposed by Panjabi, the active muscular subsystem can compensate for deficiencies in passive structures through increased muscle activation (Panjabi MM, 1992). Furthermore, studies in patients with LBP have demonstrated altered trunk muscle recruitment patterns characterized by greater activation of lumbar segmental muscles and increased co-contraction, which were predicted to increase spinal stability in modeling studies (van Dieen JH,Cholewicki J and Radebold A, 2003). Although antagonist-agonist or thoracic-lumbar co-contraction ratios were not assessed in the present study, the localized increase in MF activation is consistent with the conceptual framework and may represent a compensatory response to reduced flexion stiffness following IVD injury, aiming to restoring segmental stability during locomotion. Importantly, the addition of muscle-derived nociception (HS condition) significantly decreased EMG variability in the right MF and minimum amplitude in the left MF, compared with the spinal instability alone condition. Meanwhile, the spinal instability related increase in right MF EMG mean amplitude and variability was attenuated by the addition of muscle-derived nociception, but they were not fully reversed to baseline level. This pattern suggests that nociceptive input has an inhibitory effect on motor output, partially counteracting the instability-driven increases in MF activation.

While the mechanical consequences of IVD injury were limited and direction-specific (i.e. primarily affecting flexion) (Xiao F,Noort W,Lévénez J,Han J,van Dieën JH and Maas H, 2025), they may be sufficient to elicit localized adaptive responses. This is supported by findings in a porcine model, where IVD injury led to localized increases in corticomotor excitability specifically of the deep MF fibers that crossed the injured segment (Hodges PW et al., 2009), suggesting a central neural adaptation to IVD injury. In contrast, an ovine model showed that IVD injury led to reduced MF EMG responses to mechanical excitations, indicating impaired spinal reflex excitability (Colloca CJ et al., 2008). Therefore, these findings suggest that IVD injury may inhibit reflexive responses at the spinal level, but simultaneously trigger compensatory activation through cortical pathways to maintain spine stability and functional movement.

Our previous muscle-derived nociception-only model demonstrated reduced MF activation and ML variability (Xiao F,Noort W,Lévénez J,Han J,van Dieën JH and Maas H, 2025), consistent with nociception-related inhibition of motor output. In the present study, the presence of attenuated right MF mean activation and variability in the HS condition suggests that nociceptive input may have dampened the above mentioned adaptive neuromuscular response to IVD injury. These findings suggest that muscle-derived nociception and spine instability lead to opposing localized neuromuscular adaptations that partially cancel each other’s effects, while the overall motor patterns are largely preserved.

## Muscle-specific adaptations

The distinct activation patterns observed in MF and ML may reflect their respective role in trunk control. MF is primarily involved in stabilizing the spine, while ML contributes more to the lateral movement of the spine (Wada N,Akatani J,Miyajima N,Shimojo K and Kanda K, 2006). Our previous study demonstrated that mechanical instability following IVD injury was only in flexion rather than lateral bending (Xiao F,Noort W,Lévénez J,Han J,van Dieën JH and Maas H, 2025). Bilateral MF activation was simultaneous in all conditions. Therefore, activity of this muscle appears not to be related to lateral bending and the slightly increased activation in the IVD injury condition may indicate a compensatory strategy to counteract instability in flexion during locomotion. In contrast, the absence of changes in ML activation aligns with the finding that lateral bending mechanics were not altered, suggesting unchanged lateral stabilization demands.

Several limitations affect interpretation of the results of our study. Firstly, neuromuscular responses were assessed one week after needle puncture induced IVD injury. It is possible that neuromuscular adaptions require more time to develop, especially those involving central neural adaptations. Previous studies in rats have shown that sensorimotor restriction through hind limb immobilization during development can lead to long-lasting functional reorganization in the sensorimotor cortex over 8-13 weeks, even in the absence of structural damage (Delcour M et al., 2018; Delcour M et al., 2018). Thus, the present findings likely reflect early-stage neuromuscular responses. Future longitudinal investigations are needed to assess whether more pronounced or persistent neuromuscular adaptations emerge over time. Secondly, the side of hypertonic saline injection could only be confirmed in 5 out of 11 rats, limiting side-specific analyses. However, previous findings demonstrated that the effects of muscle-derived nociception on EMG amplitudes tended to be the same on both sides (Xiao F,Noort W,Lévénez J,Han J,van Dieën JH and Maas H, 2025). Additionally, locomotion was assessed during level ground trotting, a relatively low-demand task that may not sufficiently challenge trunk stability to reveal more pronounced neuromuscular responses. Previous studies have shown that quadrupeds exhibit greater hip joint moments (Prilutsky BI et al., 2011) and higher peak activation of the ML (Miro F et al., 2020) during uphill compared to level ground locomotion, suggesting that more demanding tasks may be more suitable to use in future studies.

## Conclusion

This study demonstrated that IVD injury, alone or in combination with muscle-derived nociception, had limited impact on overall trunk and pelvic motor control during locomotion. Trunk and pelvic kinematics and overall muscle activation patterns remained largely consistent across conditions, with only minor temporal adjustments, whereas localized, muscle-specific neuromuscular adaptations were observed. IVD injury resulted in increases in EMG mean activation and variability of the right MF. The addition of the muscle-derived nociception in the IVD injury condition reduced left MF minimum activation, and attenuated instability-related increases in right MF mean activation and variability, without fully reversing these changes. Together, these findings suggest that spinal instability and muscle-derived nociception exert opposing effects on local neuromuscular control, while the overall locomotor pattern remains robust.

## Acknowledgements

We would like to thank Amy Borsboom for her assistance with the measurements, Guus Baan for making the illustration in Fig.1, and Helga Haberfehlner for providing instructions on DeepLabCut.

This study received funding from the China Scholarship Council (grant number 202008310141). The authors declare that there are no financial interests or personal relationships that can have influenced the work reported in this paper. Data and program codes used in analysis are available to researchers for academic purpose per request.

## CRediT authorship contribution statement

**Fangxin Xiao**: Conceptualization, Data curation, Formal analysis, Funding acquisition, Investigation, Methodology, Software, Visualization, Writing-Original draft preparation, Writing-Reviewing and Editing. **Wendy Noort:** Investigation, Software, Data curation, Validation, Project administration. **Jia Han**: Supervision, Writing-Reviewing and Editing. **Jaap H. van Dieën**: Conceptualization, Methodology, Software, Resources, Supervision, Writing-Reviewing and Editing. **Huub Maas**: Conceptualization, Methodology, Resources, Visualization, Supervision, Project administration, Writing-Reviewing and Editing. All authors have read and agreed to the published version of the manuscript.

