## supplementary figure 1-10 for "Effects of lumbar disc injury and nociception on trunk motor control during rat locomotion"

| **A** | 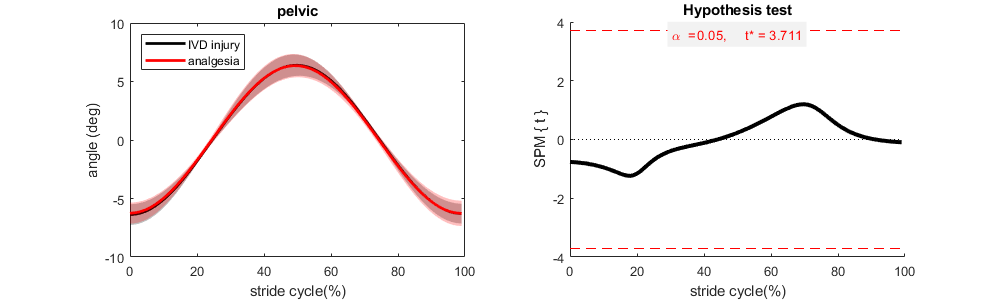 |
| --- | --- |
| **B** | 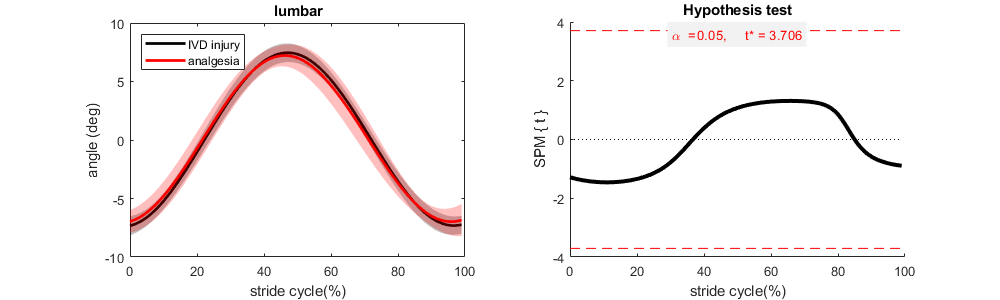 |
| **C** | 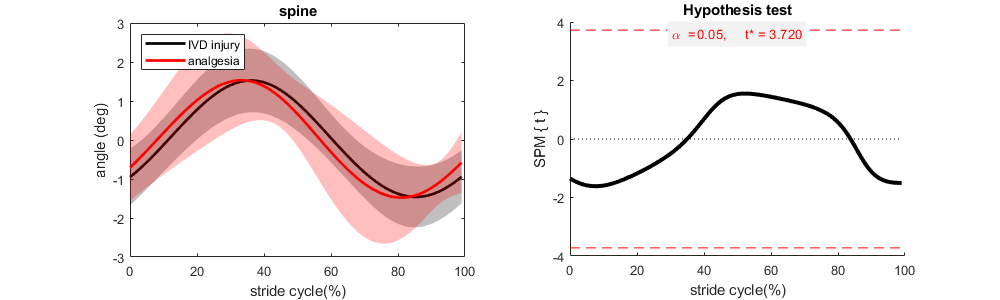 |
| **Fig.S1.** SPM analysis of the segmental angle data during locomotion for IVD injury and analgesia (carprofen injection) conditions. **(A)** pelvic, **(B)** lumbar, **(C)** spine. Joint angle data were averaged cross 10 rats and normalized to stride cycle duration. Shaded area represents mean±1SD. Treadmill speed was at 0.5m/s at all conditions. IVD, intervertebral disc injury. | |

| **A** | 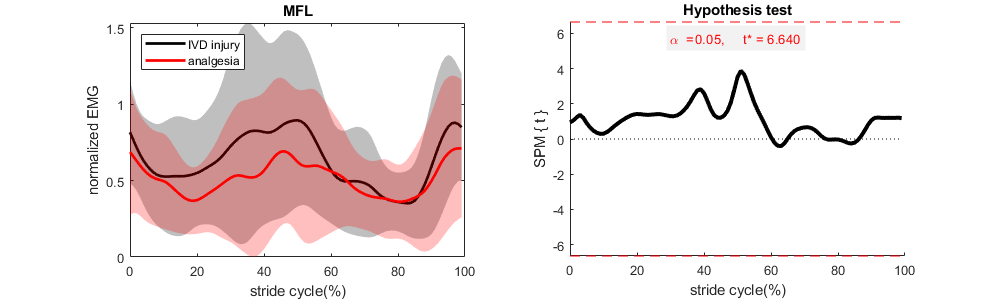 |
| --- | --- |
| **B** | 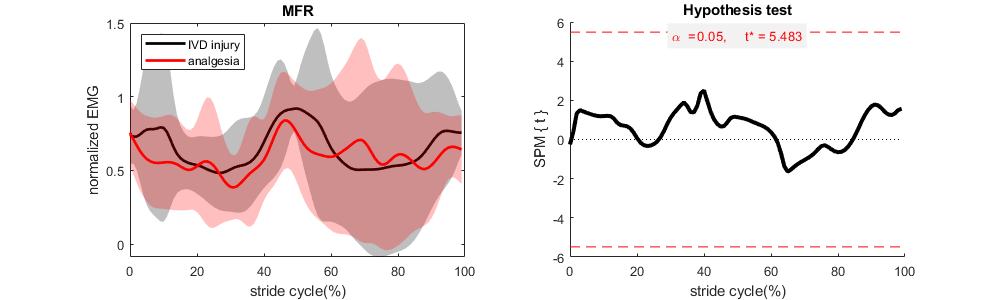 |
| **C** | 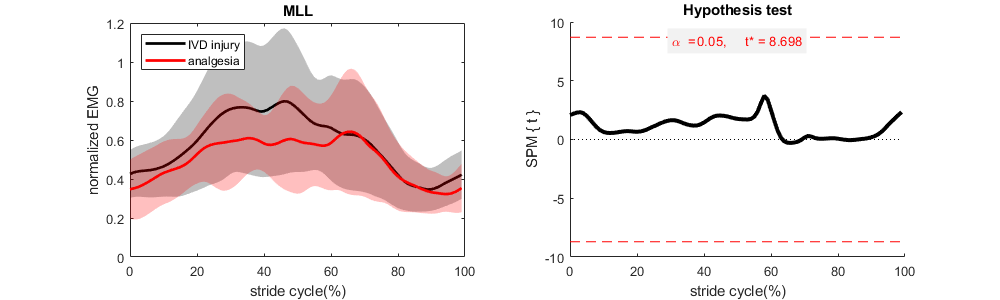 |
| **D** | 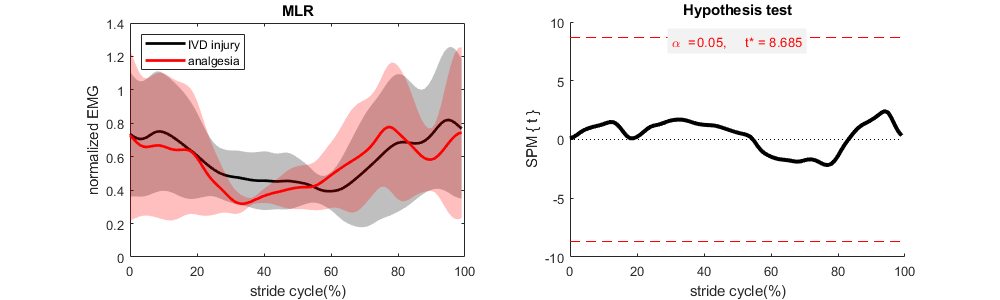 |
| **Fig.S2.** SPM analysis of the filtered rectified EMG envelope of back muscles during locomotion for IVD injury and analgesia (carprofen injection) conditions. **(A)** MFL (multifidus muscle left, n=6), **(B)** MFR (multifidus muscle right, n=8), **(C)** MLL (longissimus muscle left, n=5), **(D)** MLR (longissimus muscle right, n=5). EMG data were normalized to the stride cycle duration and peak amplitude measured during baseline. IVD, intervertebral disc injury. | |

| **A** | 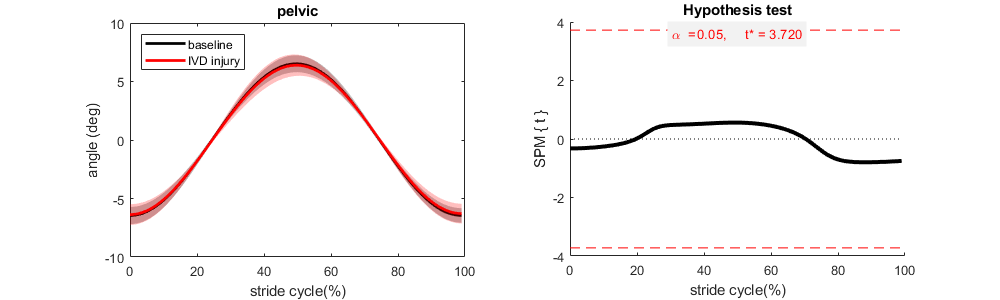 |
| --- | --- |
| **B** | 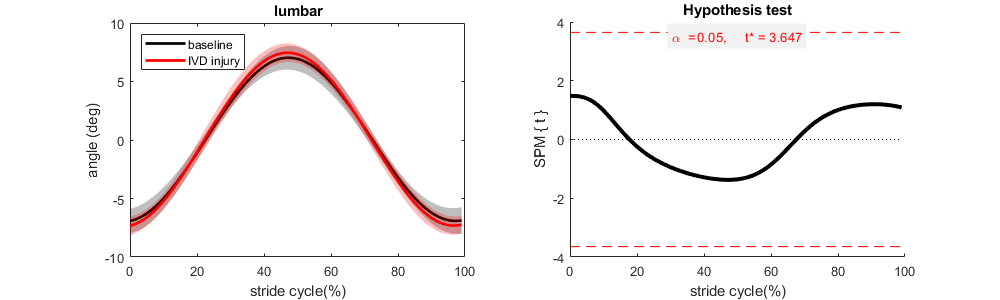 |
| **C** | 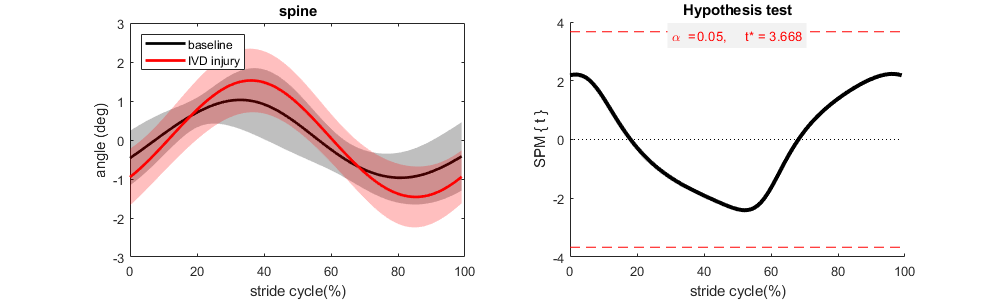 |
| **Fig.S3.** SPM analysis of the segmental angle data during locomotion for baseline and IVD injury conditions. **(A)** pelvic, **(B)** lumbar, **(C)** spine. Joint angle data were averaged cross 10 rats and normalized to stride cycle duration. Shaded area represents mean±1SD. Treadmill speed was at 0.5m/s at all conditions. IVD, intervertebral disc injury. | |

| **A** | 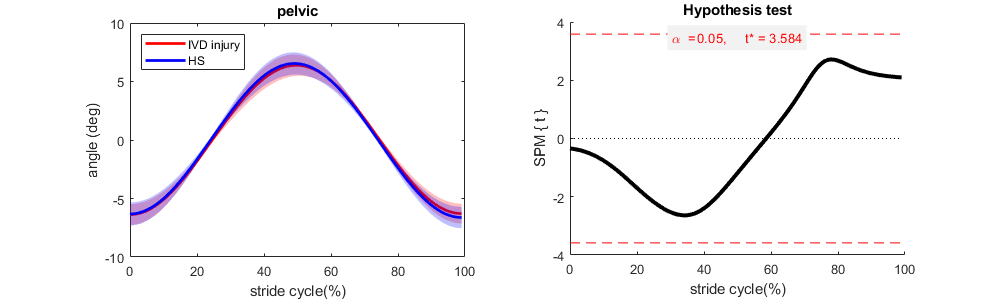 |
| --- | --- |
| **B** | 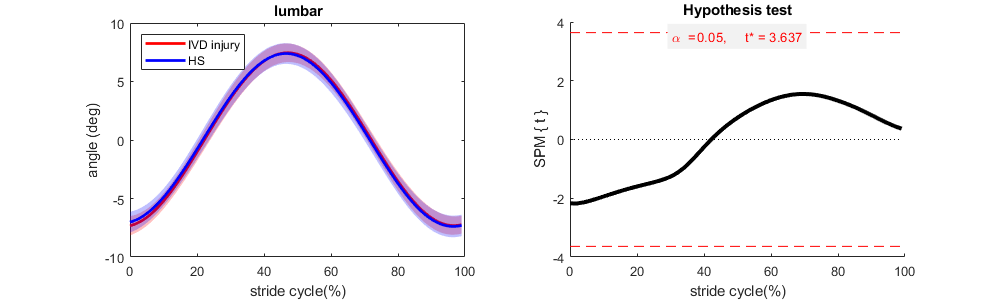 |
| **C** | 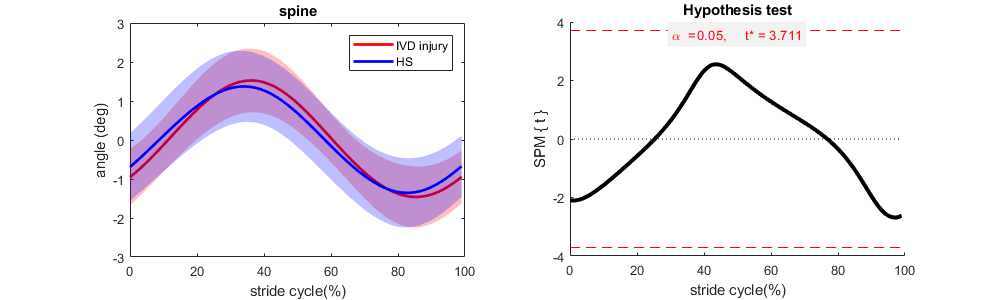 |
| **Fig.S4.** SPM analysis of the segmental angle data during locomotion for IVD injury and IVD injury + hypertonic saline injection (HS) conditions. **(A)** pelvic, **(B)** lumbar, **(C)** spine. Joint angle data were averaged cross 10 rats and normalized to stride cycle duration. Shaded area represents mean±1SD. Treadmill speed was at 0.5m/s at all conditions. IVD, intervertebral disc injury. | |

| **A** | 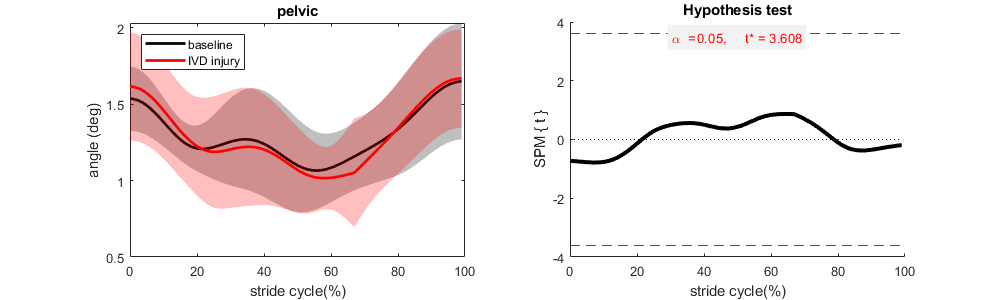 |
| --- | --- |
| **B** | 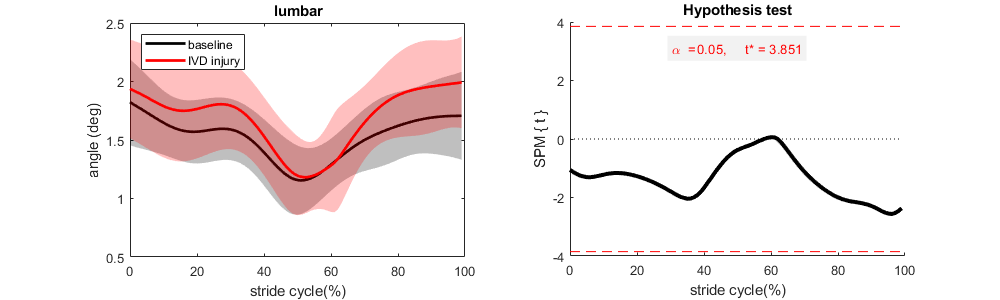 |
| **C** | 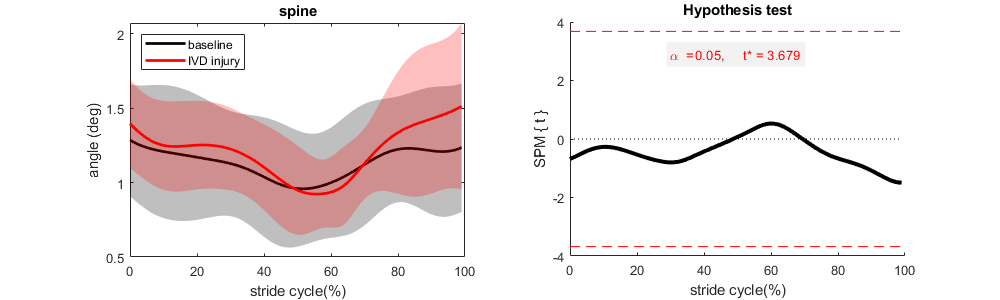 |
| **Fig.S5.** SPM analysis of the segmental angle variability during locomotion for baseline and IVD injury conditions. **(A)** pelvic, **(B)** lumbar, **(C)** spine. Joint angle variability data were averaged cross 10 rats and normalized to stride cycle duration. Shaded area represents mean±1SD. Treadmill speed was at 0.5m/s at all conditions. IVD, intervertebral disc injury. | |

| **A** | 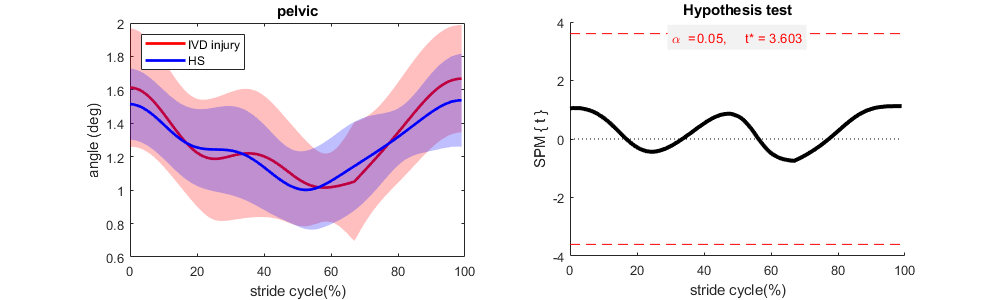 |
| --- | --- |
| **B** | 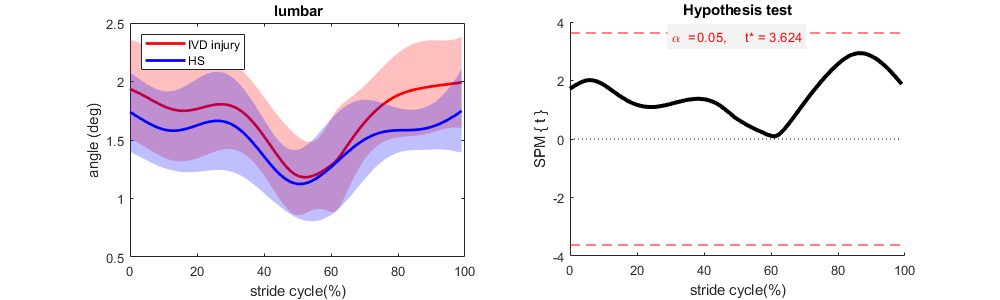 |
| **C** | 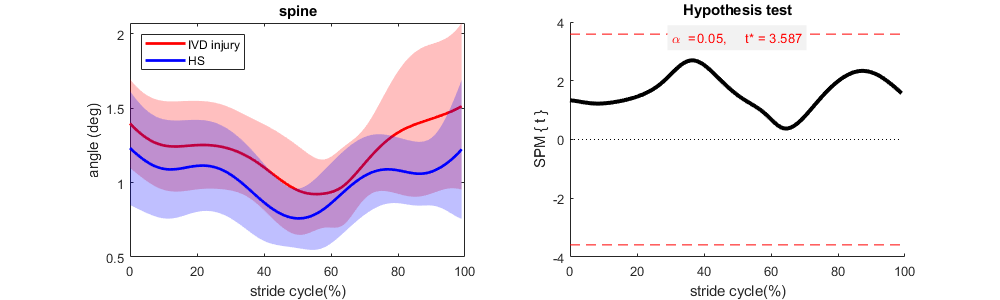 |
| **Fig.S6.** SPM analysis of the segmental angle variability during locomotion for IVD injury and IVD injury + hypertonic saline injection (HS) conditions. **(A)** pelvic, **(B)** lumbar, **(C)** spine. Joint angle variability data were averaged cross 10 rats and normalized to stride cycle duration. Shaded area represents mean±1SD. Treadmill speed was at 0.5m/s at all conditions. IVD, intervertebral disc injury. | |

| **A** | 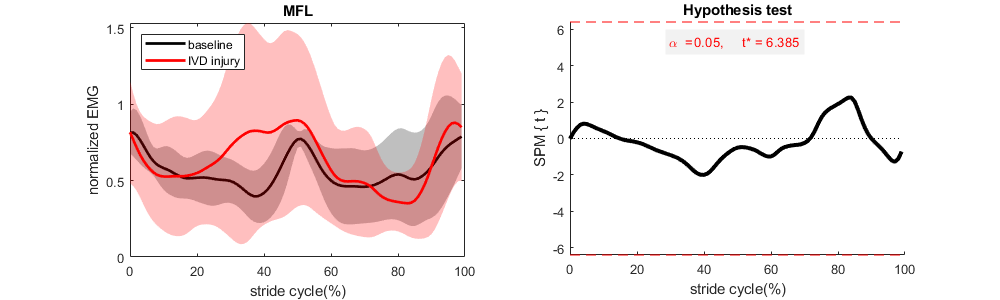 |
| --- | --- |
| **B** | 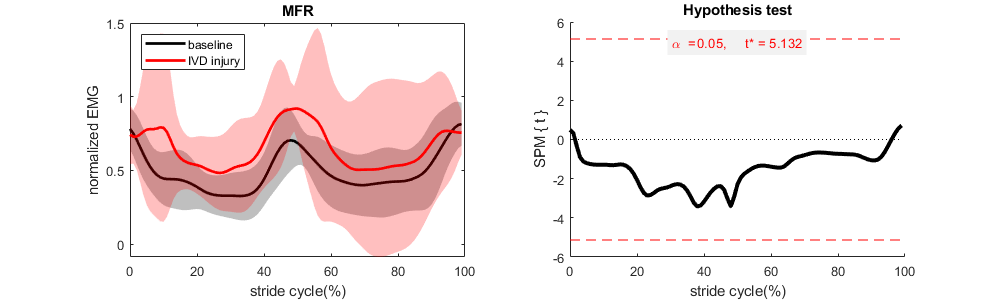 |
| **C** | 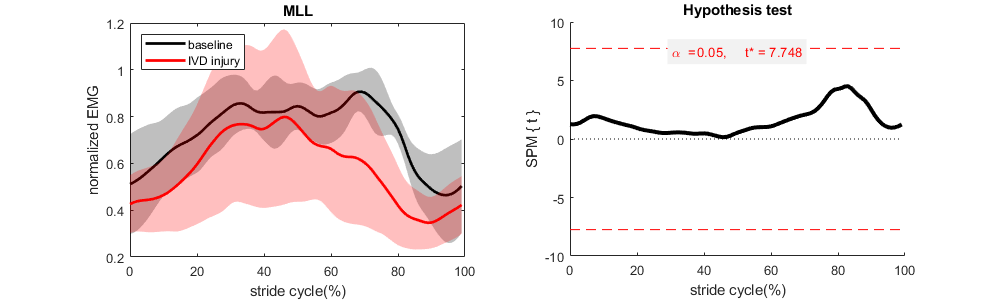 |
| **D** | 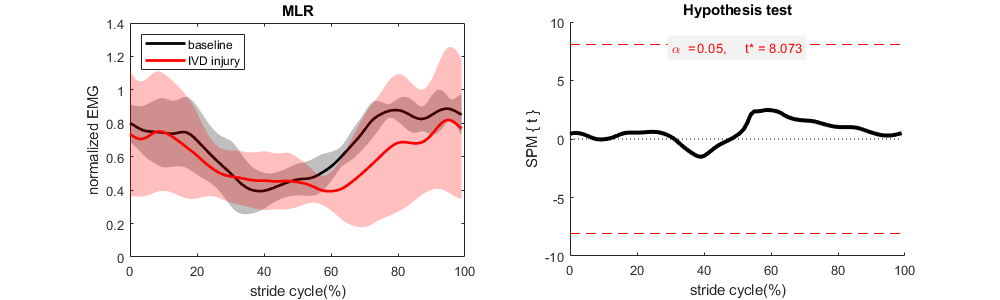 |
| **Fig.S7.** SPM analysis of the filtered rectified EMG envelope of back muscles during locomotion for baseline and IVD injury conditions. **(A)** MFL (multifidus muscle left, n=6), **(B)** MFR (multifidus muscle right, n=8), **(C)** MLL (longissimus muscle left, n=5), **(D)** MLR (longissimus muscle right, n=5). EMG data were normalized to the stride cycle duration and peak amplitude measured during baseline. IVD, intervertebral disc injury. | |

| **A** | 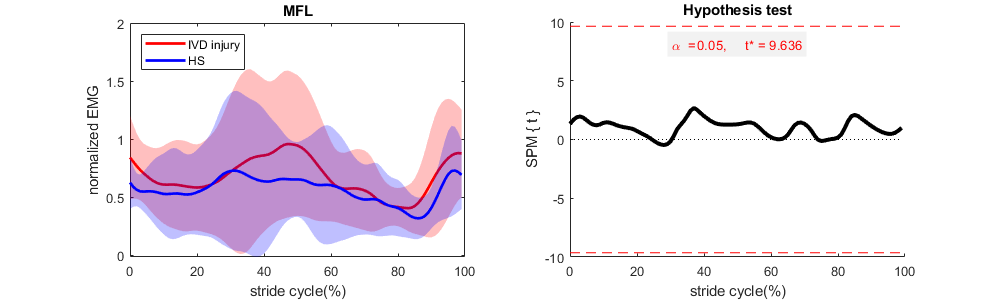 |
| --- | --- |
| **B** | 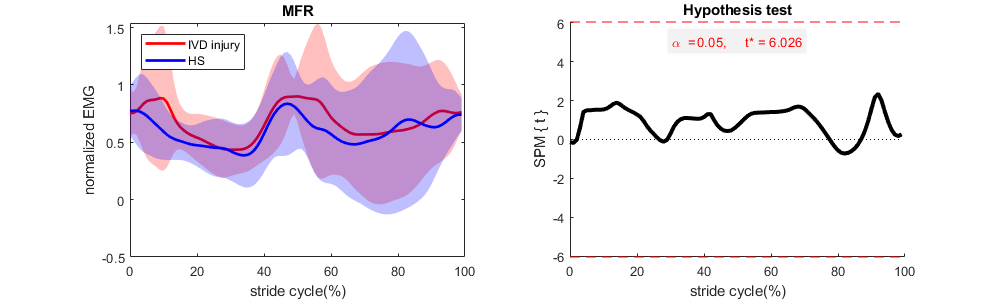 |
| **C** | 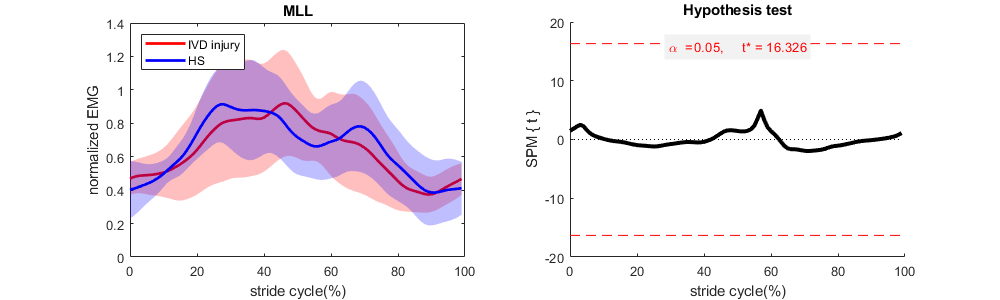 |
| **D** | 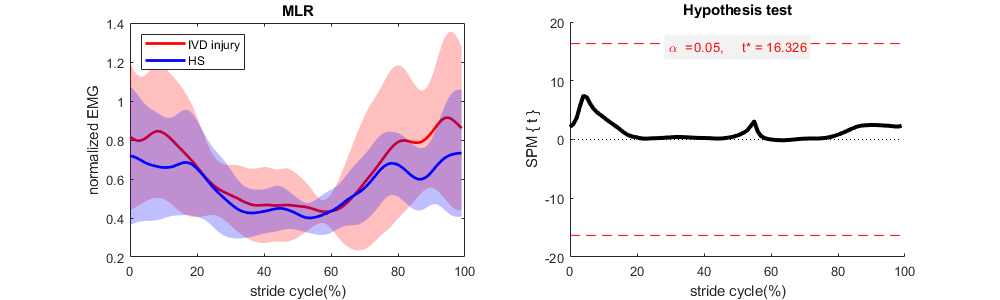 |
| **Fig.S8.** SPM analysis of the filtered rectified EMG envelope of back muscles during locomotion for IVD injury and IVD injury + hypertonic saline injection (HS) conditions. **(A)** MFL (multifidus muscle left, n=5), **(B)** MFR (multifidus muscle right, n=7), **(C)** MLL (longissimus muscle left, n=4), **(D)** MLR (longissimus muscle right, n=4). EMG data were normalized to the stride cycle duration and peak amplitude measured during baseline. IVD, intervertebral disc injury. | |

| **A** | 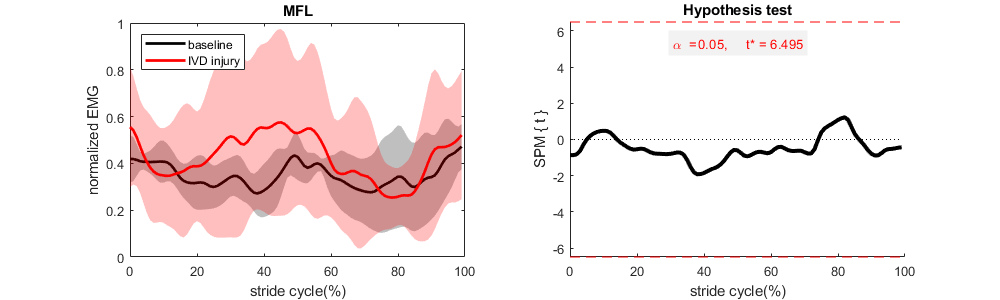 |
| --- | --- |
| **B** | 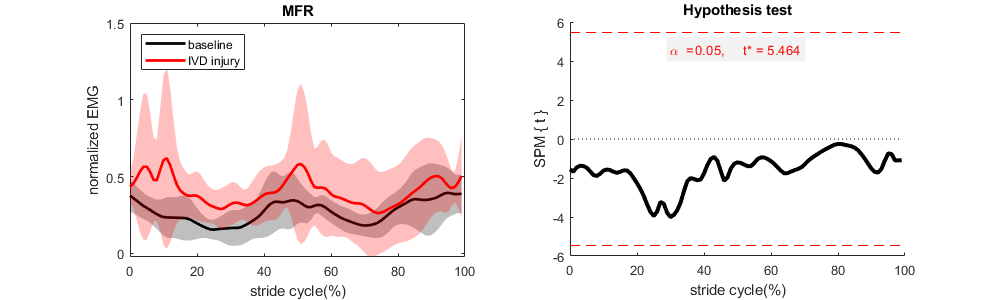 |
| **C** | 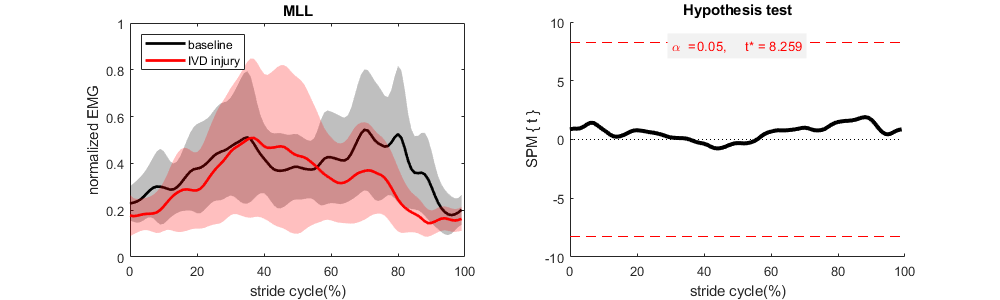 |
| **D** | 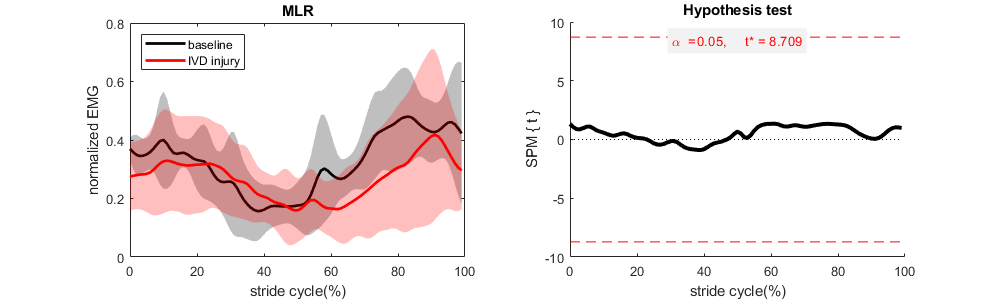 |
| **Fig.S9.** SPM analysis of the EMG variability of back muscles during locomotion for baseline and IVD injury conditions. **(A)** MFL (multifidus muscle left, n=6), **(B)** MFR (multifidus muscle right, n=8), **(C)** MLL (longissimus muscle left, n=5), **(D)** MLR (longissimus muscle right, n=5). EMG data were normalized to the stride cycle duration and peak amplitude measured during baseline. IVD, intervertebral disc injury. | |

| **A** | 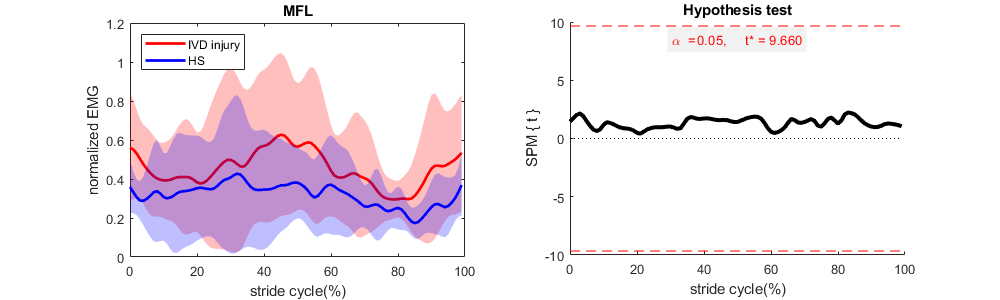 |
| --- | --- |
| **B** | 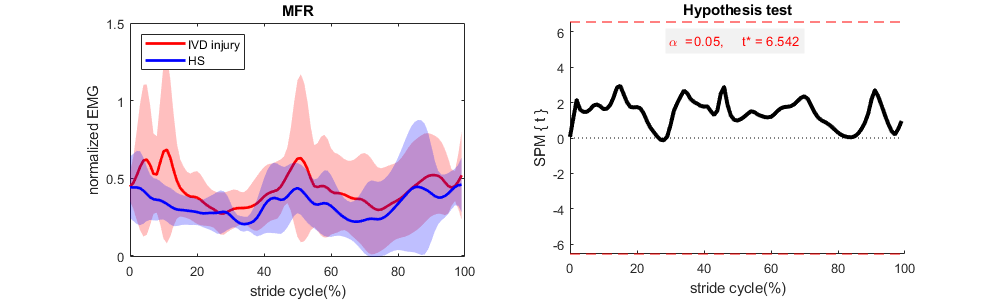 |
| **C** | 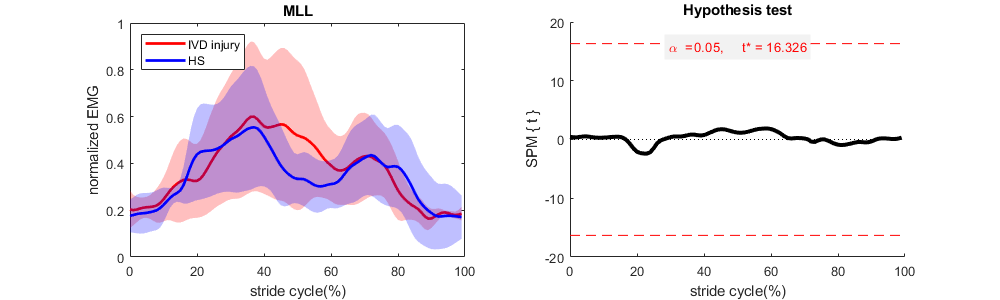 |
| **D** | 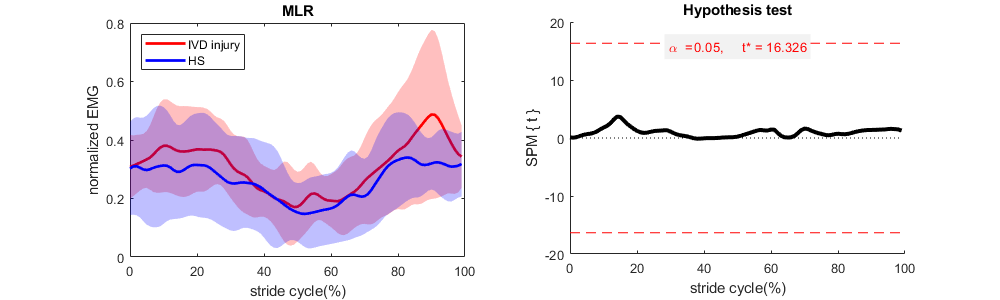 |
| **Fig.S10.** SPM analysis of the EMG variability of back muscles during locomotion for IVD injury and IVD injury + hypertonic saline injection (HS) conditions. **(A)** MFL (multifidus muscle left, n=5), **(B)** MFR (multifidus muscle right, n=7), **(C)** MLL (longissimus muscle left, n=4), **(D)** MLR (longissimus muscle right, n=4). EMG data were normalized to the stride cycle duration and peak amplitude measured during baseline. IVD, intervertebral disc injury. | |
